# Muscle activation strategies of the vastus lateralis according to sex

**DOI:** 10.1101/2021.11.01.466761

**Authors:** Yuxiao Guo, Eleanor J. Jones, Thomas B. Inns, Isabel A. Ely, Daniel W. Stashuk, Daniel J. Wilkinson, Kenneth Smith, Jessica Piasecki, Bethan E. Phillips, Philip J. Atherton, Mathew Piasecki

## Abstract

**Aim:** Despite men exhibiting greater muscle strength and fatigibility than women, it remains unclear if there are sex-based differences in muscle recruitment strategies e.g. motor unit (MU) recruitment and modulation of firing rate (FR) at normalised forces and during progressive increases in force.

**Methods:** Twenty-nine healthy male and thirty-one healthy female participants (18-35 years) were studied. Intramuscular electromyography was used to record individual motor unit potentials (MUPs) and near fibre MUPs from the vastus lateralis (VL) during 10% and 25% maximum isometric voluntary contractions (MVC), and spike-triggered averaging was used to obtain motor unit number estimates (MUNE) of the VL. Multilevel mixed-effects linear regression models were used to investigate the effects of sex at each contraction level.

**Results:** Men exhibited greater muscle strength (*p*<0.001) and size (*p*<0.001) than women, with no difference in force steadiness at 10% or 25% MVC. Women had smaller MUs and higher FR at 10% MVC (both *p*<0.02), similar to that at 25% MVC in MU size (*p*=0.062) and FR (*p*=0.031). However, both sexes showed similar increases in MU size and FR when moving from low-to mid-level contractions. There were no sex differences in any near fibre MUP parameters or in MUNE.

**Conclusion:** **I**n the vastus lateralis, women produce muscle force via different neuromuscular recruitment strategies to men which is characterised by smaller MUs discharging at higher rates. However, similar strategies are employed to increase force production from low to moderate contractions. These findings of similar proportional increases between sexes support the use of mixed sex cohorts in studies of this nature.

**Key points:**

- Increases in muscle force production are mediated by motor unit (MU) recruitment, and MU firing rate (FR).
- Women are underrepresented in studies of human neuromuscular research and markedly differ to men in a number of aspects of neuromuscular function, yet little is known of the recruitment strategies of each.
- Here we demonstrate men and women have similar vastus lateralis MU number estimates, yet women recruit smaller MUs with higher FR than men at normalised contraction levels. However, increases in force are achieved via similar trajectories of MU recruitment and MU FR in men and women.
- Although men and women exhibit divergent neuromuscular recruitment strategies to achieve normalised forces, increases in force are achived similarly and support the inclusion of mixed sex cohorts in studies of this nature.

## Introduction

Skeletal muscle contraction is regulated by central and peripheral motor and sensory nerve function and excitation-contraction coupling of muscle fibres. The fundamental neuromuscular element regulating muscle contraction is the motor unit (MU), consisting of a motor neuron and the muscle fibres it innervates ^1^. Increases in muscle force are largely mediated by two neuromuscular recruitment strategies, the recruitment of additional, progressively larger MUs, and an increase in MU firing rate (FR), referred to as MU recruitment and rate modulation, respectively ^2^. Several studies have highlighted adaptative remodelling of MUs structure and function in response to exercise training, ageing and disease ^3-6^, which influences recruitment strategies, however the majority of data are only available in men.

Men generally possess greater muscle strength than women in upper and lower extremities, which is largely explained by greater muscle size ^7^. Conversely, although task-specific, women are generally more resistant to neuromuscular fatigue when assessed at a normalised contraction level ^8^, which in the knee-extensors, is likely explained by differing fibre type ratios with a 7-23% greater proportion of type I fibres in vastus lateralis (VL) in women ^9,10^. Sex differences of the hormonal milieu also influence neuromuscular function; testosterone and estrogen are the major sex hormones in males and females, respectively, and each exhibites a range of neuroprotective effects in motoneurons, such as dendritic maintenance and axonal sprouting ^11^. Furthermore, hormonal metabolites are associated with the release of brain-derived neurotrophic factors (BDNF) ^12^, which are key mediators of synaptic plasiticity ^13^. Acutely, differences in sex hormones partly explain the variability in fatigability in women across phases of the menstrual cycle ^14^. Such differences in the hormonal milieu are difficult to experimentally control for and may explain why women are often underrepresented in studies of neuromuscular physiology ^15^.

Surface electromyography (EMG) has been commonly applied to study sex-based differences of neuromuscular function and muscle recruitment strategies ^16,17^. However, such approaches are limited by the distance between activated MUs and recording electrodes ^18^, offering poor MU yield in women, and in some cases, being influenced by adjacent muscles ^19^. These limitations can be overcome with the use of intramuscular EMG (iEMG), which also has the added benefit of revealing further electrophysiological parameters relevant to MU size and complexity ^20^. Although we have previously reported the sex-based divergent trajectory of MU FR from middle to older age in long-term trained master athletes ^21^, comparisons of normative values in healthy young men and women at differing contraction levels are unknown. The aims of the present study were to compare individual MU properties and muscle recruitment strategies, as well as the MU number estimates (MUNEs) in the VL of healthy young men and women. We hypothesised several parameters would differ at normalised contraction levels, with no sex-based differences in recruitment strategies when moving from a low-to a mid-level contraction.

## Materials and Methods

### Ethics approval

This research was approved by the University of Nottingham Faculty of Medicine and Health Sciences Research Ethics Committee (C16122016, 160-0121, 186-1812, 103-1809, 302-1903) and was conducted between 2019 and 2021 in accordance with the Declaration of Helsinki.

### Participants

Twenty-nine healthy male and thirty-one healthy female participants, aged 18-35 years, were recruited via advertisement posters in the local community. All the participants volunteered to take part in the studies and provided written informed consent. Prior to enrolment, all participants completed a comprehensive clinical examination and metabolic screening were conducted at the School of Medicine, Royal Derby Hospital Centre. All articipants were recreationally active. Participants with metabolic disease, lower limb musculoskeletal abnormalities, acute cerebrovascular or cardiovascular disease, active malignancy, uncontrolled hypertension, or those on medications that impact muscle protein metabolism or modulate vascular tone were excluded.

### Anthropometry

Body mass and height were measured using calibrated scales and a stadiometery, respectively for the calculation of body mass index (BMI). Ultrasound was used to measure the cross-sectional area of the VL using an ultrasound probe ((LA523 probe, B-mode, frequency range 26–32 Hz, and MyLabTM50 scanner, Esaote, Genoa, Italy) at the anatomical mid-point of the muscle which was identified between the greater trochanter and the mid-point of the patella with participants lying supine. Ultrasound images were acquired aligning the superior edge of the probe following a middle-to-lateral direction position on the skin, beginning, and ending the image capture at aponeurosis borders. A water-based conductive gel was applied on the surface of the ultrasound probe to enhance the fidelity of the image without causing excessive contact pressure on the skin during the acquisition of the images. Images were subsequently analysed using ImageJ software (National Institutes of Health, USA) to quantify CSA measurements. The mean area of three images was taken as CSA. The CSA of eight female participants was measured using magnetic resonance imaging (MRI) with a T1-weighted turbo 3D sequence on a 0.25-T G-Scan (Esaote, Genoa, Italy). Continuous transversal images with a 6-mm slice were acquired and analysed by using Osirix imaging software (Osirix medical imaging, Osirix, Atlanta, GA, United States) through tracing around the VL following the contour of the aponeurosis. VL CSA values are available for 23 men and 19 women.

### Muscle strength and force steadiness

The maximum isometric voluntary contraction force (MVC) of the right knee extensor was assessed with the participants sitting in a custom-built chair with hips and knees flexed at ∼90 degrees. The lower leg was securely attached to a force dynamometer with non-compliant straps (purpose-built calibrated strain gauge, RS125 Components Ltd, Corby, UK) slightly above the medial malleolus. A seat belt was fastened across the pelvis to avoid superfluous movement of the upper trunk during the test. To obtain the external knee joint moment arm, the distance from centre of the force strap to the lateral femoral condyle was measured. After a standardised warm-up of submaximal contractions, participants were instructed to perform a maximal isometric contraction with real-time visual feedback and verbal encouragement. This was repeated a further two times, with 60 second rest intervals between each, and the highest value was accepted as MVC. Peak torque during the selected MVC was also determined. After determination of the MVC, participants were instructed to perform between four and six sustained isometric contractions at 10% and 25% MVC, respectively, each lasting 12-15 seconds with a target line displayed on the screen in front of the participants (with iEMG, below, Figure 1). Participants had 20-30 seconds rest between each contraction. Prior to the assessment, participants were allowed a single familiarisation practice at each contraction level. Force steadiness was quantified as the coefficient of variation of the force [CoV; (SD/mean) × 100]. The mean CoV at each contraction level was calculated from the middle two contractions.

**Figure 1.**
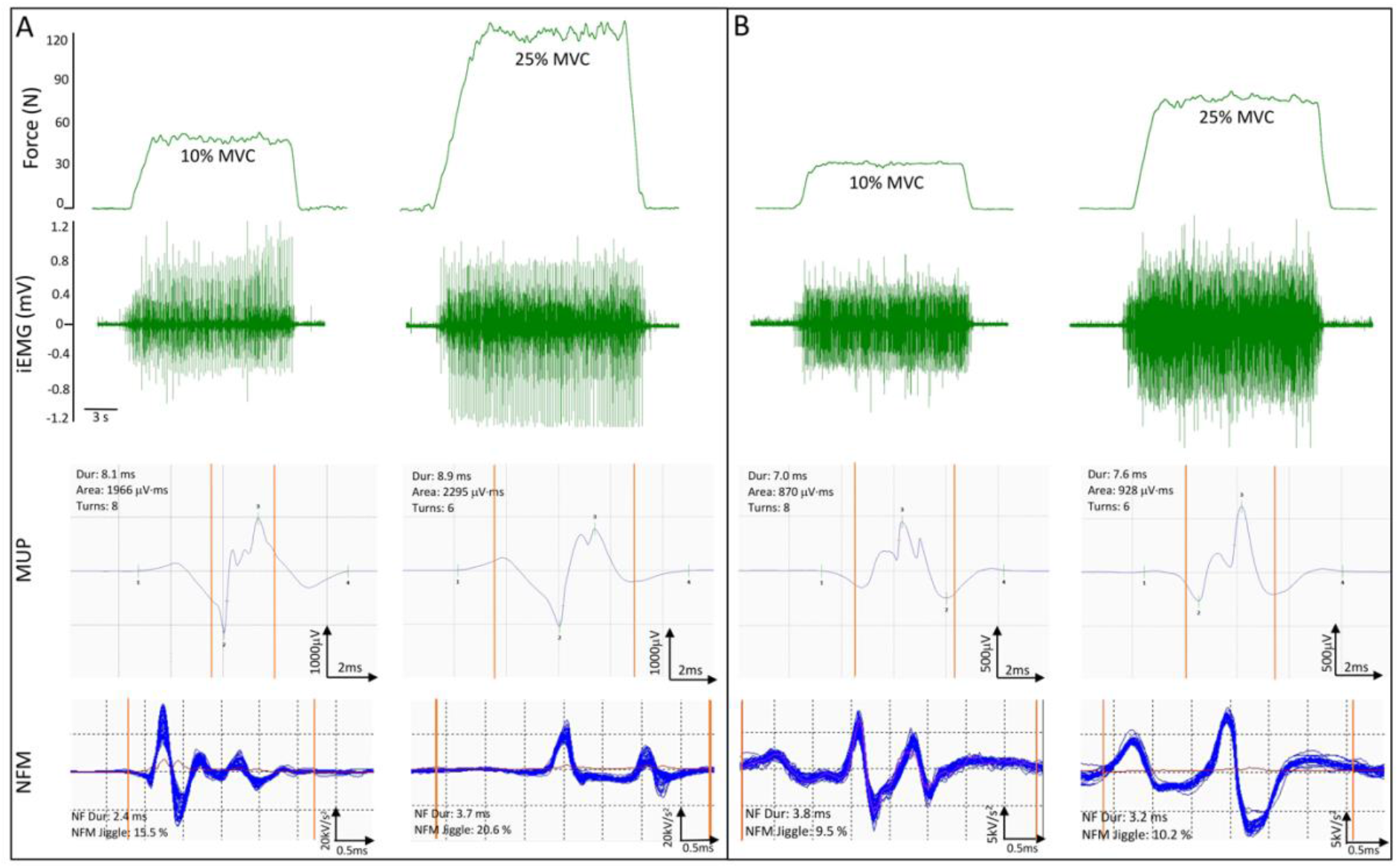
Representative data from a male (A) and a female (B) participant. Top panels show knee extensor force traces at 10% and 25% of MVC, and corresponding intramuscular electromyography (iEMG) raw data recorded from the vastus lateralis. A representative MUP template and corresponding NFM shimmer plot isolated from each contraction are shown below each iEMG signal. Vertical orange lines on MUPs and NFMs indicate the start and end time of the NFM. Abbreviations: N, newtons; mV, millivolt; MVC, maximum voluntary contraction; MUP; motor unit potential; Dur, duration; NF, near fibre; NFM, near fibre MUP; kV, kilovolt; μV, microvolt; ms, millisecond.

### Surface electromyography (sEMG)

An active recording sEMG electrode (disposable self-adhering Ag-AgCl electrodes; 95 mm^2^, Ambu Neuroline, Baltorpbakken, Ballerup, Denmark) was placed over the motor point located around the mid-point of the VL, identified using a cathode probe (Medserve, Daventry, UK) to apply percutaneous electrical stimulation at 400 V, pulse width of 50 μs and current of around 8 mA (DS7A Digitimer, Welwyn Garden City, Hertfordshire, UK) with a self-adhesive anode electrode (Dermatrode, Farmadomo, NL) placed over the right gluteus. A reference electrode was placed over the patella tendon and a common ground electrode placed over the patella. The common ground electrode served for both sEMG and iEMG measurements. sEMG signals were sampled at 10kHz, and bandpass filtered between 5 and 5 kHz (1902 amplifier, Cambridge Electronics Design Ltd., Cambridge, UK) and digitized with a CED Micro 1401 data acquisition unit (Cambridge Electronic Design) for offline analysis.

### Compound muscle action potential (CMAP)

The CMAP of the VL was evoked by a manually triggered stimulator (model DS7A; Digitimer) using percutaneous stimulation (Medserve, Daventry, UK) of the proximal femoral nerve (approximately halfway between the anterior superior iliac spine and the pubic tubercle) with a carbon-rubber anode electrode (Dermatrode self-adhering electrode, 5.08 cm in diameter; Farmadomo Linde Homecare Benelux Bv, Leiden, The Netherlands) placed over the skin overlying the gluteus muscle. The stimulator voltage was fixed at 400 V and the pulse width at 50 μS, with the current increased incrementally until the M-wave amplitude plateaued. At this point, the current was increased again by ∼30 mA to ensure supramaximal stimulation, ensuring a sharp rise time of the negative peak of the m-wave.

### Intramuscular electromyography (iEMG)

A 25-mm disposable concentric needle electrode (N53153; Teca, Hawthorne, New York, USA) was inserted at the muscle belly of VL, adjacent to the recording surface electrode over the motor point, to a depth of 1.5-2 cm depending on the muscle size. The iEMG shared the same ground electrode as the sEMG, which was placed over the patella. iEMG signals were recorded using Spike2 (Version 9.06), sampled at 50 kHz and bandpass filtered at 10 Hz to 10 kHz (1902 amplifier; Cambridge Electronic Design Ltd, Cambridge, UK) and stored for future off-line analysis.

Prior to EMG and CMAP assessments, participants performed a series of voluntary, low-level contractions once the needle was positioned to ensure adequate signal to noise ratio, thus ensuring the recording needle electrode was close to depolarizing fibres. Each participant then performed the sustained voluntary isometric contractions as detailed above (Figure 1). After a 10% and 25% MVC contraction, to avoid repeat sampling of the same MUs, the needle electrode was repositioned by the combinations of twisting the bevel edge 180 degrees and withdrawing by ∼5 mm. This process was repeated until four to six contractions from spatially distinct areas (from deep to superficial portions) were recorded ^18^. Participants had ∼30 seconds rest between each contraction.

### EMG analysis

Decomposition-based quantitative electromyography (DQEMG) software was used to detect motor unit potentials (MUPs), extract motor unit potential trains (MUPTs) generated by individual MUs from the iEMG signals and estimate, via ensemble averaging, their corresponding surface MUPs (sMUPs) from the sEMG signals ^22^. MUPTs that were composed of MUPs from more than one MU or had fewer than 40 MUPs were excluded. The occurrence times of individual MUPs within a MUPT were used to trigger and align sEMG signal epochs for ensemble-averaging to produce an estimate of their corresponding sMUPs. The sMUPs estimated from signals recorded during 25% MVC contractions were use to represent MU size. All MUP and sMUP templates were visually inspected and their markers adjusted, where required, to correspond to the onset, onset of negative phase (sMUP only), end, and positive and negative peaks of the waveforms.

MUP amplitude was measured from the maximal positive and negative peaks and the MUP area was taken as the total area within the MUP duration (onset to end) and is indicative of MU size. The number of phases and turns are measures of MUP complexity and are classified as the number of components above or below the baseline (phases) and a change in waveform direction of at least 25 μV (turns), which indicates the level of temporal dispersion across individual muscle fibre contributions to a single MUP. MU FR was assessed as the rate of MUP occurrences within a MUPT, expressed as the number of occurrences per second (Hz). A near fibre MUP (NFM) is defined as the acceleration of its corresponding MUP (Figure 1) and calculated by applying a second-order, low-pass differentiator to the MUP which effectively reduces the recording area of the needle electrode to within ∼350 μm, thereby ensuring only potentials from fibres closest to the needle electrode significantly contribute to the NFM and reducing interference from distant active fibres of other MUs. NFM jiggle is a measure of the shape variability of consecutive NFMs of an MUPT expressed as a percentage of the total NFM area. NFM segment jitter is a measure of the temporal variability of individual fibre contributions to the NFMs of a MUPT. It is calculated as a weighted average of the absolute values of the temporal offsets between matched NFM segments of consecutive isolated (i.e., not contaminated by the activity of other MUs) NFMs across an MUPT expressed in microseconds. NFM dispersion is the time, in ms, between the first and last MU fibre contributions ^20^.

### Motor unit number estimates (MUNE)

The MUNE value was derived by dividing the negative peak area of the ensemble averaged mean surface MUP (msMUP) from 25% MVC into the negative peak area of the CMAP ^23^. A msMUP is an ensemble average of the negative-peak onset aligned, sMUPs of the MUs sampled from a muscle. The negative peak area of the msMUP was divided into the negative peak area of the electrically evoked CMAP ^24^. MUNE values are available for 15 men and 15 women.

### Statistical analysis

All of the statistical analysis was performed using RStudio (Version 1.3.959 for macOS) ^25^. Descriptive statistics of participant characteristics are presented as *mean* ± *standard deviation (SD). Student’s unpaired t-test* was used to compare physical parameters (age, BMI, MVC, CSA, and force steadiness). As multiple MUs were recorded from each participant, *multi-level mixed-effect linear regression analysis* was performed to investigate these MU parameters with sex and contraction level as factors through the package *lme4* (Version 1.1.23) ^26^. In the linear mixed models, the first level was single motor unit; single motor units were clustered according to each participant to form the second level, which was defined as the participant level. This linear mixed-effect modelling framework is suitable for data of this nature as it: i) incorporates the whole sample of extracted MUs not just the mean values obtained from each participant, which preserves variability within and across participants simultaneously to the greatest extent; ii) handles missing data better than an *analysis of variances (ANOVA)* framework as the removal of a single missing observation has a much smaller effect in the mixed model; and iii) provides coefficient estimates that indicate the magnitude and direction of the effects of interest ^27^. Interactions were first examined and where not present they were removed from the model, sex and contraction level were explored individually. The results are displayed as coefficient estimates, 95% confidence intervals, and *p*-values. Standardized estimates were calculated through the package *effectsize* (Version 0.4.5) ^28^ for forest plotting. For data visualisation, individual participant means and group means were shown in box-and-jitter plots. Statistical significance was assumed when *p* < 0.05. Based on the models used, *p* values close to 0.05 were also addressed ^29^.

## Results

The means and standard deviations for the participant’s characteristics are shown in Table 1. Significant differences between men and women were detected for weight, height, BMI, peak torque and VL CSA (all *p*<0.05). There were no significant sex differences for age and force steadiness (both *p*>0.05). Individual values are shown in Figure 2.

**Table 1.** Participant characteristics.

| Measure | Men (n=29) | Women (n=31) | P value |
| --- | --- | --- | --- |
| Age (years) | 23.7(5.0) | 22.9(3.6) | 0.490 |
| Weight (kg) | 81.2(11.6) | 63.1(10.1) | **<0.001** |
| Height (cm) | 180.3(7.5) | 165.8(6.8) | **<0.001** |
| BMI (kg/m<sup>2</sup>) | 25.0(2.9) | 23.0(3.2) | **0.012** |
| Peak Torque (Nm) | 251.65(67.72) | 158.95(46.69) | **<0.001** |
| VL CSA (cm<sup>2</sup>) | 28.35(6.43) | 19.00(3.90) | **<0.001** |
| Force CoV-10% | 4.51(1.21) | 4.98(1.18) | 0.139 |
| Force CoV-25% | 3.36(1.17) | 3.24(0.74) | 0.638 |

Data are reported as mean (standard deviation). Values in bold reflect statistically significant ( $p<0.05$ ) results. Abbreviations: BMI, body mass index; VL CSA, vastus lateralis cross-sectional area.

**Figure 2.**
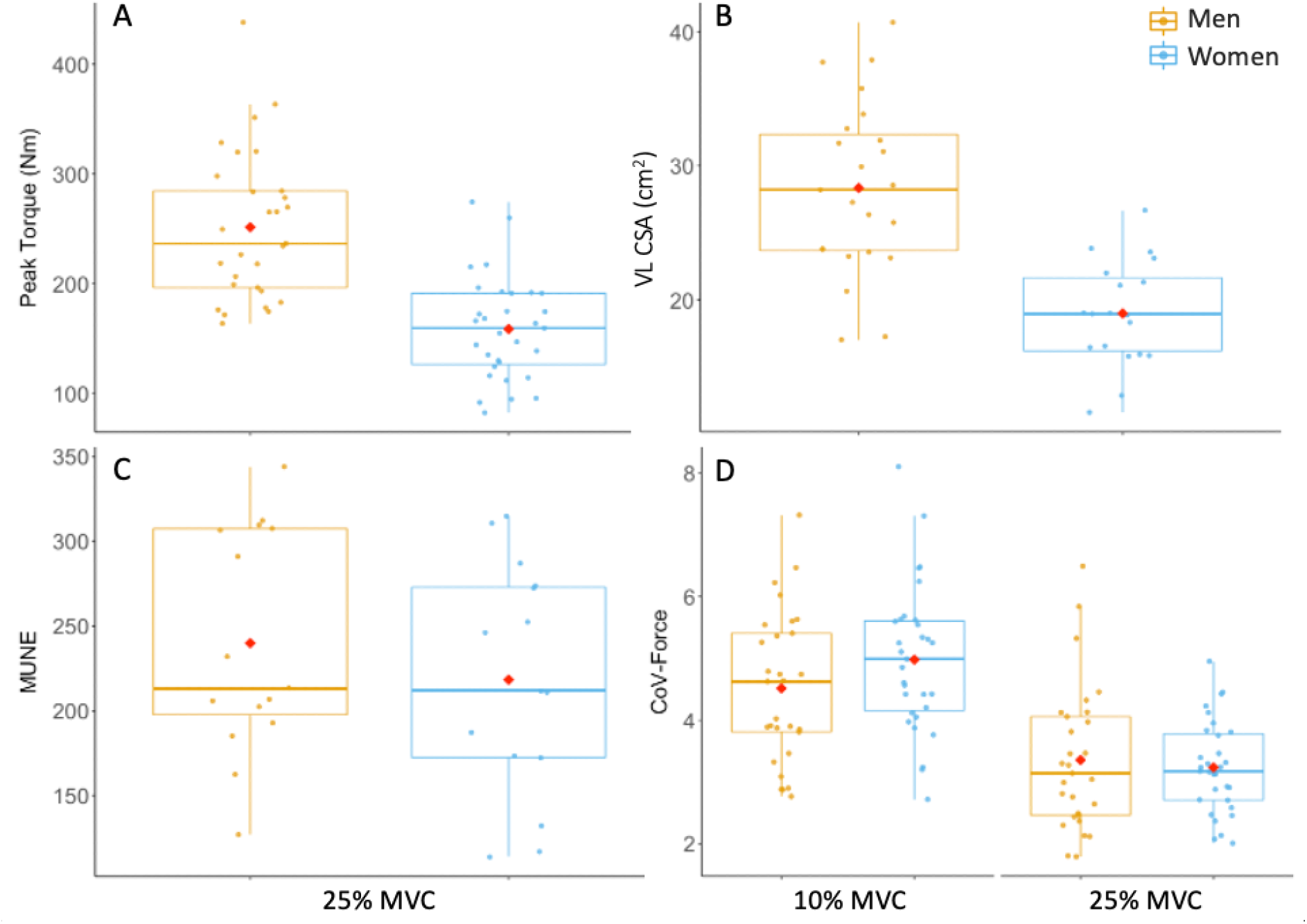
Box-and-jitter plots of the individual participant means, and group means (red dot) of **(A)** peak torque, **(B)** vastus lateralis cross-sectional area, **(C)** motor unit number estimates (MUNE), and **(D)** force steadiness at 10% and 25% maximum voluntary contraction (MVC), in men (yellow) and women (blue). Abbreviations: CoV, coefficient of variation.

A total of 1645 MUs were analysed in men and 1207 in women. At 10% MVC, the mean number of MUs isolated per person was 26 in men and 17 in women; at 25% MVC, the mean number of MUs isolated per person was 31 in men and 22 in women. Individual mean values for all functional, MU and NFM parameters are sown in Figures 2-5. There were no significant interactions between sex and contraction level in any of the MU parameters. When interactions were removed from the model, multilevel linear regression revealed women had greater MU FR at both 10% (mean; M: 8.08 Hz; W: 8.79 Hz) and 25% (M: 8.62 Hz; W: 9.20 Hz) MVC (both *p*<0.05, Table 2, Figure 3A). This was matched by a non-significant trend (*p*<0.10) for greater MU FR vaiability in women at both contraction levels (Table 2, Figure 3B). MUP duration was shorter at 10% (M: 8.37 ms; W: 6.61 ms) and 25% (M: 8.24 ms; W: 6.84 ms) MVC in women when compared to men (both *p*<0.01, Table 2, Figure 4E). MUP area was smaller in women at 10% (M: 741μV·ms; W: 531 μV·ms) (*p*=0.006), with a non-significant trend at 25% MVC (M: 1005 μV ·ms; W:775μV &ms) (*p*=0.062). There were no significant sex-based differences in any other MU characteristic (Table 2).

**Table 2.**
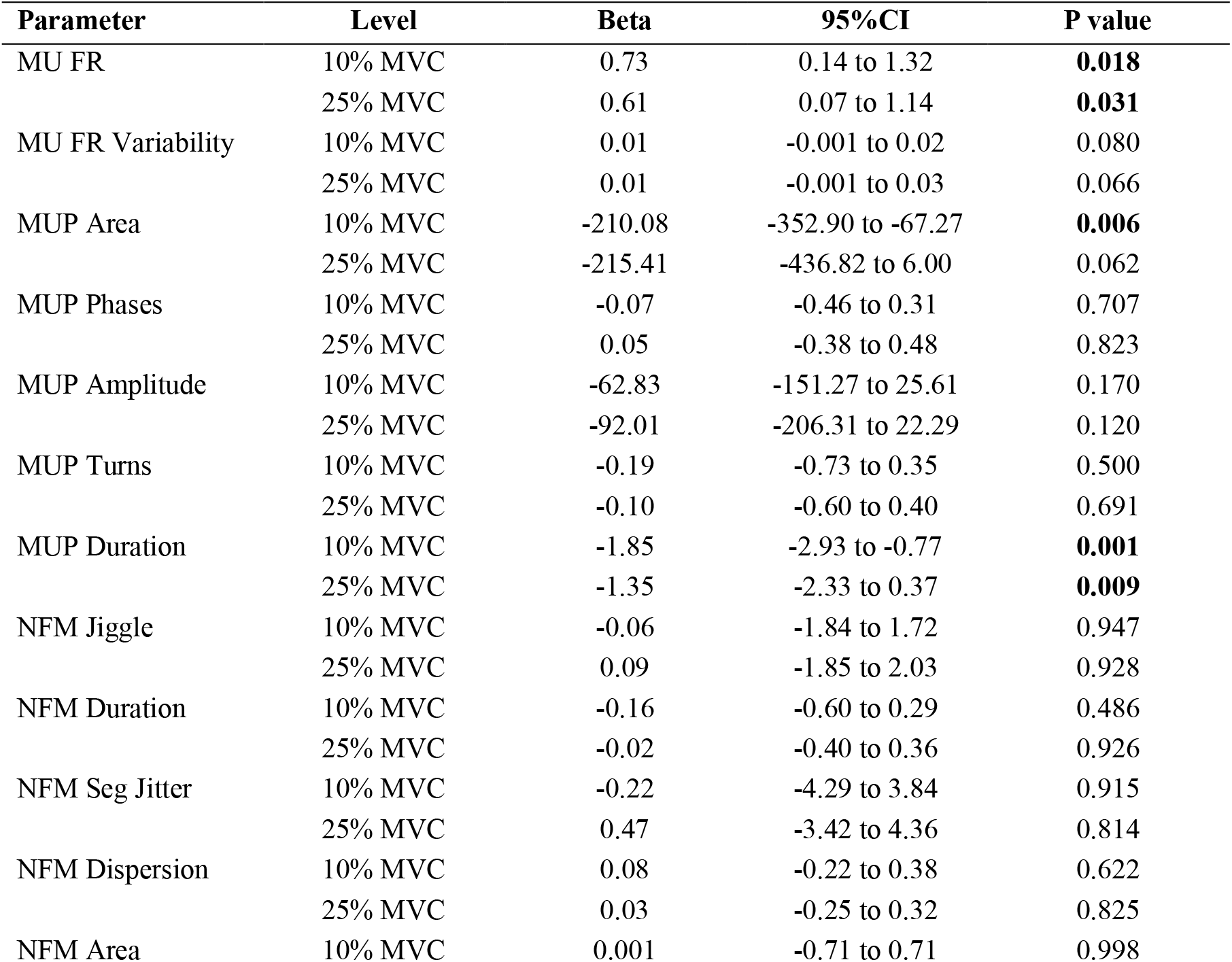
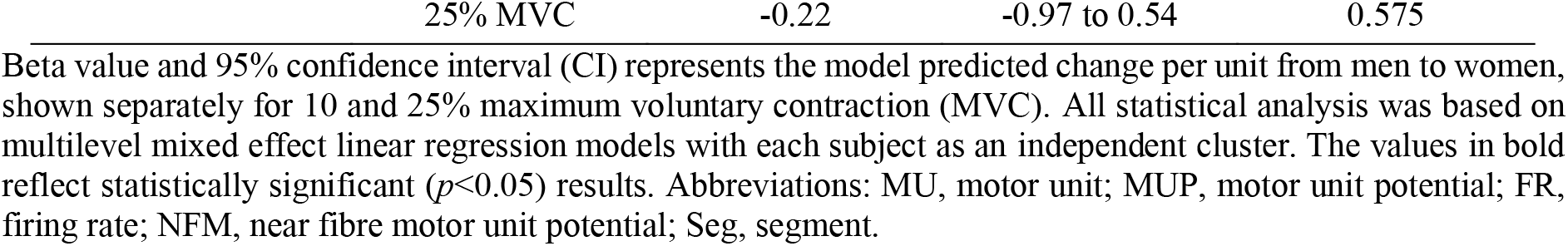
Motor unit properties in different sexes

**Figure 3.**
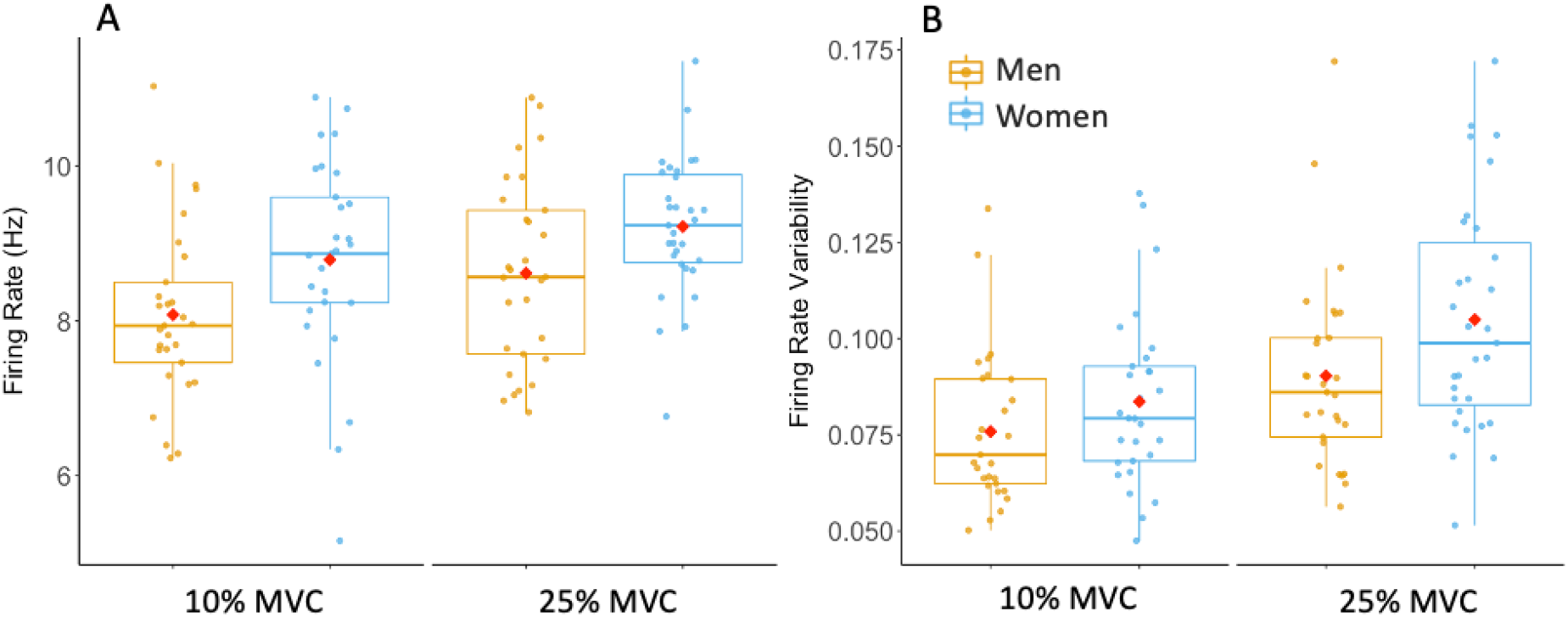
Box-and-jitter plots of the individual participant means, and group means (red dot) of **(A)**motor unit (MU) firing rate and **(B)** firing rate variability in men (yellow) and women (blue) at 10 and 25% maximum voluntary contraction (MVC).

**Figure 4.**
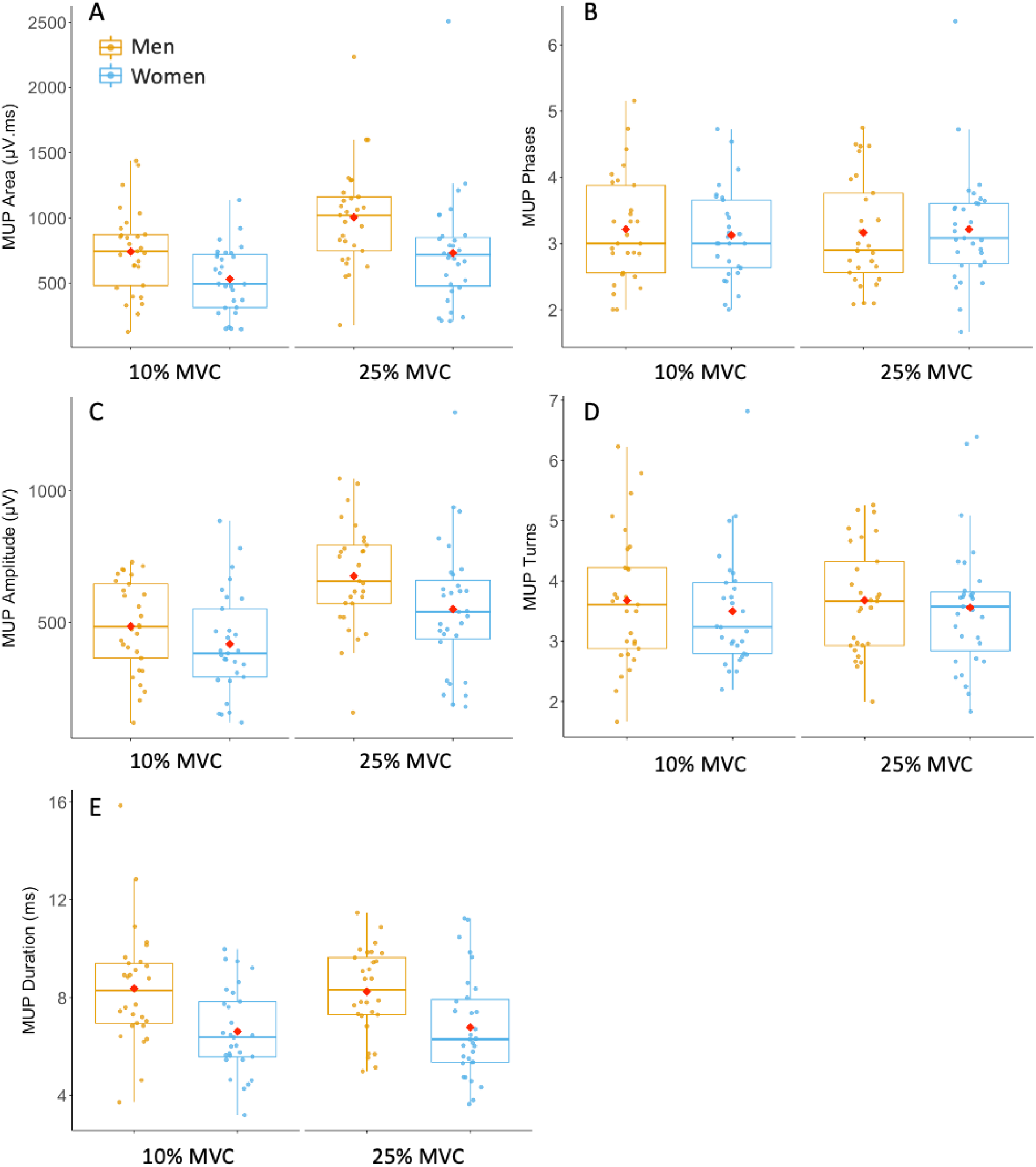
Box-and-jitter plots of the individual participant means, and group means (red dot) of motor unit potential (MUP) properties in men (yellow) and women (blue) at 10 and 25% maximum voluntary contraction (MVC).

**Figure 5.**
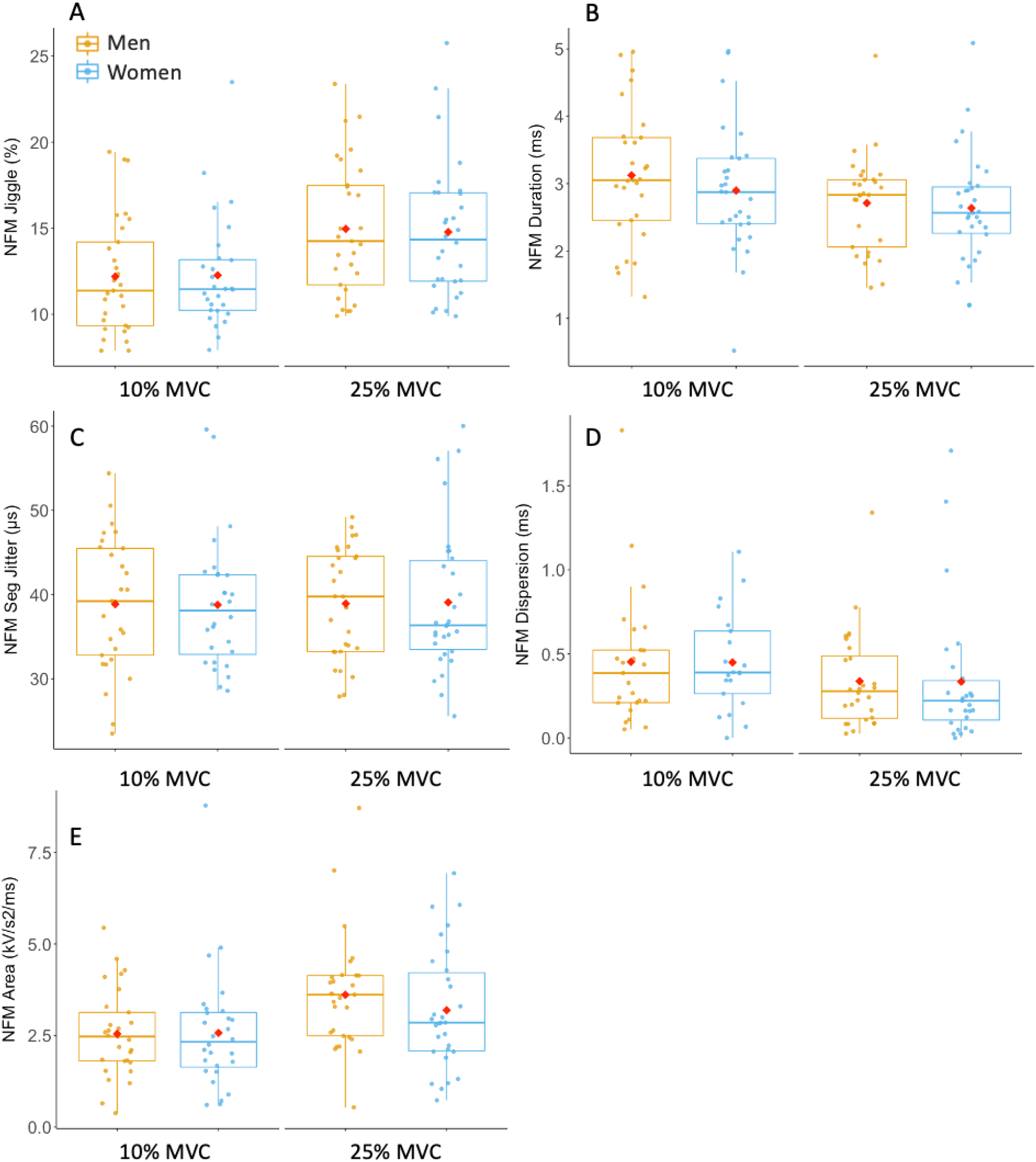
Box-and-jitter plots of the individual participant means, and group means (red dot) of near fibre motor unit potential (NFM) properties in men (yellow) and women (blue) at 10 and 25% maximum voluntary contraction (MVC).

With increasing contraction level, both men and women exhibited higher MU FR and MU FR variability, as well as greater MUP amplitude, and larger MUP area (all *p*<0.001, Table 3, Figures 3 and 4). NFM segment jitter, NFM duration and NFM area were also larger with the higher contraction level, increasing to a similar extent in men and women (all *p*<0.001, Table 3, Figure 6). There were no interactions between sex and contraction level in any of the MU parameters, indicating the difference from 10 to 25% MVC did not differ between men and women (Figure 6).

**Table 3.** Motor unit properties at different contraction levels

| Parameter | Sex | Beta | 95%CI | P value |
| --- | --- | --- | --- | --- |
| MU FR | Men | 0.50 | 0.28 to 0.71 | **<0.001** |
|  | Women | 0.43 | 0.18 to 0.69 | **0.001** |
| MU FR Variability | Men | 0.02 | 0.01 to 0.02 | **<0.001** |
|  | Women | 0.02 | 0.01 to 0.02 | **<0.001** |
| MUP Area | Men | 273.62 | 215.81 to 331.43 | **<0.001** |
|  | Women | 199.35 | 148.13 to 250.56 | **<0.001** |
| MUP Phases | Men | -0.05 | -0.14 to 0.04 | 0.302 |
|  | Women | 0.10 | -0.02 to 0.21 | 0.098 |
| MUP Amplitude | Men | 188.21 | 153.75 to 222.67 | **<0.001** |
|  | Women | 142.40 | 106.27 to 178.53 | **<0.001** |
| MUP Turns | Men | 0.05 | -0.11 to 0.20 | 0.565 |
|  | Women | 0.09 | -0.08 to 0.26 | 0.316 |
| MUP Duration | Men | -0.16 | -0.52 to 0.20 | 0.381 |
|  | Women | 0.13 | -0.16 to 0.42 | 0.371 |
| NFM Jiggle | Men | 3.08 | 2.38 to 3.79 | **<0.001** |
|  | Women | 3.08 | 2.34 to 3.82 | **<0.001** |
| NFM Duration | Men | -0.40 | -0.54 to -0.26 | **<0.001** |
|  | Women | -0.34 | -0.50 to -0.18 | **<0.001** |
| NFM Seg Jitter | Men | 0.77 | -0.37 to 1.91 | 0.188 |
|  | Women | 0.51 | -0.85 to 1.87 | 0.463 |
| NFM Dispersion | Men | -0.09 | -0.23 to 0.05 | 0.228 |
|  | Women | -0.05 | -0.28 to 0.18 | 0.655 |
| NFM Area | Men | 0.98 | 0.68 to 1.29 | **<0.001** |
|  | Women | 0.72 | 0.39 to 1.05 | **<0.001** |

Beta value and 95% confidence interval (CI) represents the model predicted change per unit from 10 to 25% maximum voluntary contraction (MVC), shown separately for men and women. All statistical analysis was based
on multilevel mixed effect linear regression models with each subject as an independent cluster. The values in bold reflect statistically significant ( $p<0.05$ ) results. Abbreviations: MU, motor unit; MUP, motor unit potential; FR, firing rate; NFM, near fibre motor unit potential; Seg, segment.

**Figure 6.**
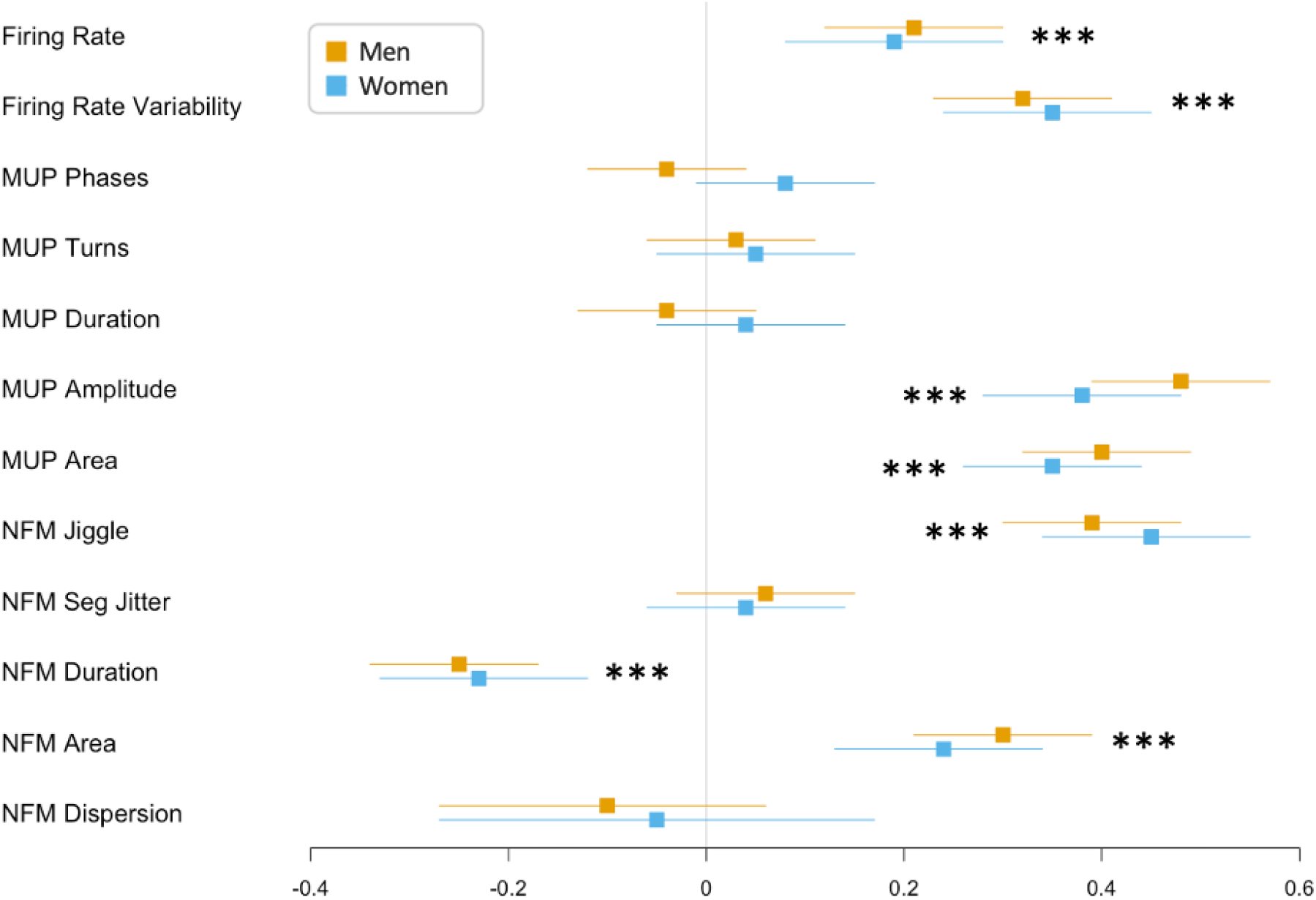
Forest plots of the standardised regression coefficient estimate for associations between motor unit characteristics and contraction levels in men and women models. Beta value and 95% confidence intervals (CI) represents the standardised model predicted change per unit from 10 to 25% maximum voluntary contraction (MVC). All statistical analysis was based on multilevel mixed effect linear regression models with each subject as an independent cluster, largely maintaining the motor unit variability within each subject. Standardised values of each parameter make the comparisons executable between men and women. *** =*p*<0.001.

## Discussion

This is the first study to compare muscle recruitment strategies and motor unit number estimates of the VL using iEMG techniques in healthy young men and women. Despite men having stronger and larger muscles, there were no differences in force steadiness at either lower or moderate contraction levels between sexes. At each contraction level assessed, women displayed smaller markers of MU size and greater MU FR, indicating differing recruitment strategies to achieve a normalised force. When assessing the difference between contraction levels, both men and women exhibited higher MU FR and greater MUP size, which differed to a similar extent in both sexes, indicating a similar recruitment strategy to generate proportional increases in force. In addition, there was no significant sex-based differences in motor unit number estimates of the VL. These data reveal divergent neuromuscular recruitment strategies between sexes to achieve a normlised force, which follow similar trajectories with increasing force.

Consistent with previous studies, women exhibited 33% smaller muscle size (CSA of the VL), which was reflected in a 31% lower strength ^7,30,31^. The greater MU FR of women shown here in VL is in an agreement with some but not all previously published data and again highlights probable muscle specific confounders. For instance, women exhibited higher MU FR and MU FR variability compared with men in elbow flexors, flexor digitorum indicis, biceps, knee extensors and tibialis anterior ^32-35^. However, others reported no sex difference in knee extensors during 30% MVC ^36^ and significantly greater MU FR at 100% MVC in tibialis anterior in men ^37,38^. These differences in intrinsic motoneuron excitability may be explained by sex-specific levels of persistent inward currents, which amplify and prolong synaptic input to MUs and affected by the levels of central monoanmines ^39^, which are reported to be higher in women ^40^. As expected, in the current study MU FR increased with increasing force levels to a similar extent in both men and women, accompanied by greater MU FR variability. Recruitment threshold dominates at lower force levels, whereas MU FR is more significant at higher forces ^41,42^, which may explain the increased variability with increased force. Although differeng at each contraction level, the similar proportional increase in MU FR in men and women indicates both sexes follow similar discharge pattern increases from low to mid-level contractions.

Despite large differences in muscle strength, force steadiness - representing the ability to hold a constant force, which is also influenced by MU FR and its variability ^32,43-45^, did not differ between sexes at either contraction level. Differing from the current findings, Inglis ^35^ found that women had a greater MU FR variability and greater fluctuation in steadiness than men during dorsiflexion in tibiais anterior muscles, which may indicate a muscle specific sex difference. In the current study, both men and women exhibited greater force steadiness at 25% MVC compared 10% MVC, consistent with Inglis’s finding that very high and low force outputs have greater fluctuations compared to moderate force outputs ^35,46^.

The size of a MU can be estimated by the size of the MUP recorded using intramuscular electrodes. As previously mentioned, men typically exhibit larger muscle size than women, ^9,47^, with increases in force mediated by recruitment of additional larger MUs and increases in MU FR. Here MUP area and duration were smaller in women which reflects smaller MU size. When viewed alongside the greater MU FR in women, it suggests that at the normalised force levels assessed here, women are more reliant on MU FR than on recruitment of larger MUs, when compared to men. As expected, markers of MU size increased at larger contraction levels, as larger MUs are recruited to produce larger forces. Again, the trajectory of each was similar for men and women indicating MU recruitment strategies moving between these force levels do not differ between sexes.

A NFM is derived from a MUP, such that is primarily composed of contributions from MU fibres close to the intramuscular electrode ^20^. Here there were no sex differences in any NFM parameters at either contraction level. When comparing 10% MVC and 25% MVC contractions, NFM area increased, while NFM duration decreased, albeit to a similar extent in both sexes. These contraction induced alterations may be the result of the activation of larger MU fibres with greater conduction velocity during higher level contractions.

Increases in NFM instability, as measured by NFM jiggle or NFM segment jitter can reflect increases in neuromuscular junction (NMJ) transmission instability with age ^21,48-50^ and in diabetic neuropathy ^3^. In the current study NFM instability, as measured by NFM jiggle, increased with contraction level for both sexes, and to similar extents. NFM jiggle is based on variability in the amplitudes of NFM shapes, and although these amplitude changes are normalized by the size of the NFM, it is possible that these increases with contraction level may be due to the recruitment of larger MUs with more MU fibres contributing to larger NFMs at 25% MVC. Combined with the lack of a sex difference in NFM segment jitter, it is clear NMJ transmission instability in the VL is sensitive to contraction level and is similar in heathy young men and women. However, there were no statistically significant contraction-based differences in NFM segment jitter, which is based on variability in the occurrence times of NFM segments and is not affected by NFM size, indicating it is less sensitive to the influences of contraction level.

The mean values of MUNE in men (240 ± 66) and women (218 ± 68) reported here are similar to those we have previously reported in male cohorts ^5,50^, and highlights the repeatability of this method in this muscle group when applying identical experimental procedures. Although the MUNE should be viewed as an index relative to the number of MUs within a muscle and not a true anatomical count, the similar values reported here in men and women support minimal sex-based differences. Combined with the small difference in MU size at 25% MVC, which is lower than total muscle size difference, the current data support the notion that sex-based differences in total muscle size are largely explained by greater individual fibre size in men ^47^.

Although providing a high level of detail of MU structure and function via MUPs and NFMs sampled in deep and superficial muscle regions, regardless of subcutaneous tissue amount, iEMG is sensitive to contraction level and reliably identifying individual MU activity at high levels in this muscle can be problematic. Therefore, data presented here were obtained during low and mid-level contractions only. Secondly, we did not control for hormonal fluctuations in women naturally occurring during the menstrual cycle, and although there is limited evidence to confirm this, these fluctuations may influence several parameters included here. This is a direct comparison of MU features in young men and women, and further investigations concerning neural drive and influence of hormones on neural drive are still required to further understand the sex-based differences in the motor nervous system.

In summary, when compared to men, women exhibited smaller VL MUs which exhibited higher MU FR, when assessed at a single normalised contraction level. However, both men and women showed similar increases in MU size and MU FR from a low-to a mid-level contraction, indicating a similar neuromuscular recruitment strategies. These results suggest that although sex-based neuromuscular differences are apparent at a single contraction level, relative differences between levels are similar in this widely studied muscle group. These data do not support the notion of excluding women from studies of this nature.

## Acknowledgements

We thank all of the participants for their enthusiastic involvement in this study. We are grateful to Mr Daniel McCormick for assistance with data collection.

## Funding

This work was supported by the Medical Research Council [grant number MR/P021220/1] as part of the MRC-Versus Arthritis Centre for Musculoskeletal Ageing Research awarded to the Universities of Nottingham and Birmingham, and was supported by the NIHR Nottingham Biomedical Research Centre. The views expressed are those of the author(s) and not necessarily those of the NHS, the NIHR or the Department of Health and Social Care. This work was also supported by a Physiological Society Research Grant awarded to MP.

## Author contributions

All authors contributed to the conception and design of the work. Y.G., E.J.J., T.B.I, I.A.E., J.P., and M.P. contributed to the acquisition and analysis of the data. YG analysed the data and drafted the manuscript. B.E.P, P.J.A., D.J.W, K.S., D.W.S, and M.P. provided comments. All authors have approved the final version of the submitted manuscript for publication and are accountable for all aspects of the work. All persons designated as authors qualify for authorship, and all those who qualify for authorship are listed.

## Conflict of interest

The authors have no conflict of interest to declare.

## Data availability statement

The datasets generated and analysed during the current study are available from the corresponding author upon reasonable request.

## Notes

### Competing Interest Statement

The authors have declared no competing interest.

